# Effects of muscle mass on muscle force predictions in human movement

**DOI:** 10.64898/2026.03.30.714909

**Authors:** Ing-Jeng Chen, Ameur Latreche, Stephanie A. Ross, Javier A. Almonacid, Taylor JM Dick, Evie E. Vereecke, James M. Wakeling

**Affiliations:** Department of Biomedical Physiology and Kinesiology, Simon Fraser University, Canada; Faculty of Kinesiology, University of Calgary, Canada; Department of Mathematics, Simon Fraser University, Canada; School of Biomedical Sciences, The University of Queensland, Australia; Department of Development and Regeneration, KU Leuven, Belgium

**Keywords:** skeletal muscle, muscle mechanics, muscle mass, inertia, cyclic contractions

## Abstract

Muscle mass significantly influences skeletal muscle behaviour, potentially explaining why traditional massless Hill-type models struggle to predict the forces generated by larger muscles during dynamic, submaximal contractions. However, the applicability of mass-enhanced Hill-type models in human locomotion remains unexplored. Here, we compared the predicted force from a 1D mass-enhanced Hill-type muscle model with a traditional 1D massless Hill-type muscle model across a range of experimentally measured human movements. Kinematic and electromyographic data were collected from twenty participants performing locomotor tasks and supplemented with existing cycling data. Muscle size was geometrically scaled by factors from 0.1 to 10, which causes lengths to be scaled proportionally, cross-sectional area and peak isometric force F_0_ with the square, and mass with the cube of the factor. Muscle tissue mass (inertia) and cadence increased the differences between mass-enhanced and massless predictions of force and power. At high cadence and the largest scale, the normalized root mean square difference between force traces reached 7% of F_0_, (averaged across muscles). However, differences between models were minimal (<1%) at human-sized scale 1. Real muscle additionally deforms in 3D, we still do not know the extent to which this extra dimensionality affects muscle forces for these human movements.

## Introduction

Mass is a fundamental physical quantity of all matter, including skeletal muscle. Deformation and motions of a muscle require accelerations of the muscle mass, and these result from forces acting on that mass. During a contraction, energy is required for these inertial accelerations, and is associated with the kinetic energy of the muscle: the cost of this internal energy detracts from the energy available for the muscle to perform external work (Ross et al., 2021; Wakeling et al., 2020). The presence of a muscle’s mass thus influences its contraction dynamics, including the rates of force development (Günther et al., 2012), maximum contraction speed (Böl & Reese, 2008; Ross & Wakeling, 2016), external work and power (Ross, Ryan, et al., 2018; Ross et al., 2021; Wakeling et al., 2020), and the efficiency of contraction (Ross & Wakeling, 2021). Accordingly, it is important to determine how these mass effects influence model predictions of force, work, and power in humans across a range of movements, an aim that motivates the present study.

Musculoskeletal simulations typically represent the human body as a series of rigid segments, each assigned specific mass and inertial properties. Most existing biomechanical simulations assume segments to be rigid bodies with fixed inertial properties; however, several previous studies have explicitly incorporated the mass and inertia of soft tissues (including muscle tissue) to better represent their influence on segmental dynamics and limb motion (He et al., 1991; Cole et al., 1996; Pai, 2010; Guo et al., 2020; Verheul et al., 2023). These studies showed how the inertia of soft tissue (muscle mass) affects the dynamics and motion of the body segments; however, they did not consider how muscle mass affected the contractile performance and mechanical output of the muscles per se. The current study specifically addresses this gap by examining how muscle mass influences the muscle’s own contractile properties during various movement tasks.

The influence of muscle mass on the contraction dynamics of a muscle should increase with muscle size. The inertial resistance due to the mass opposes changes in motion or rest. Activated muscle fibres must produce force to surpass this inertial resistance, accelerating their own as well as the neighboring mass of inactive fibres within the muscle. Specifically, with geometric scaling the force a muscle can generate scales with its cross-sectional area (∝ L^2^, the square of its length), but the inertial mass of the tissue that would be accelerated by this force scales with its volume (∝ L^3^, the cube of its length). Consequently, larger muscles experience higher inertial loads relative to the force they can generate. This disparity becomes even more pronounced during submaximal contractions because the total force available to accelerate the inertial loads is less, and during high-frequency contractions where the rapid changes in velocity cause high accelerations. Indeed, experiments have demonstrated a decrease in the maximum shortening velocity when muscle activation is reduced (Holt et al., 2014; Josephson & Edman, 1988), and a decrease in the internal accelerations of a muscle when internal mass is added (Ross et al., 2020). Modelling studies have predicted how increased size results in a decrease in the relative work and power that a muscle can generate during cyclic contractions (Ross, Ryan, et al., 2018; Ross et al., 2021; Ross & Wakeling, 2021). One other crucial aspect that may affect muscle mass on contraction dynamics is the wobbling mass dynamics, which arises from the serial arrangement of masses and elasticities (the muscle’s natural oscillation). Gruber et al. (1998) found that mass dynamics are crucial factors in high-impact responses (jumping, running), in which they critically shape the courses of the ground reaction force as well as the joint moments and forces. Christensen et al. (2017) used isolated muscles to show the impact of muscle mass on force generation and cross-bridge dynamics. Later, Christensen et al. (2023) described the relevant wobbling frequencies with a model with multiple point masses. Furthermore, contractile properties also change with animal size, as different myofilament isoforms are present in the muscles. Thus, for instance, the maximum contraction velocity of the muscle also changes with animal size (without considering the influence of muscle mass) (Rome et al. 1990), which also affects force generation during dynamic contractions.

These findings raise an open question about how important it is to include mass in the muscle models. So, the use of 1D mass-enhanced muscle models is a step in the right direction, as it allows the influence of muscle inertia to be investigated under defined boundary conditions.

When a muscle changes length it must change in girth as it attempts to maintain its volume, thus deformations of a muscle tissue can occur in all three dimensions. The internal energetic cost to accelerating the muscle mass will thus accumulate across all three of its dimensions during contraction (Ross et al., 2021). Three-dimensional models of muscle that account for the muscle volume, density and thus mass have shown that the mass effects for dynamic contractions average 17% in relative work and power at full activation (for muscle that is scaled to be 3.5 times longer than human size), with the effects exceeding this at submaximal activations (Ross et al., 2021).

However, one-dimensional (1D) Hill-type muscle models (Zajac, 1989; Delp et al., 2007) remain the practical standard in large-scale musculoskeletal simulations due to their computational efficiency, yet they do not inherently account for the volume, density, or mass of muscle tissue (Wakeling et al., 2023). By ignoring these parameters, traditional 1D models essentially treat large muscles and small fibers alike, assuming they share the same intrinsic properties regardless of size or mass, and thereby omit scale effects. Under this assumption, muscles with identical geometric proportions but different masses yield the same normalized force and mass-specific mechanical work. Recent 1D mass-enhanced Hill-type models (Günther et al., 2012; Ross & Wakeling, 2016, 2021) allow us to study the scale effects. These models have demonstrated that placing point masses in-series with contractile segments induces inertial forces that reduce net work output at high contraction velocities; however, these models did not quantify the effect of the muscle mass on contractile dynamics for human-sized muscles during tasks relevant to everyday locomotion. Our study extends this framework by applying both a traditional 1D *massless* Hill-type muscle model and a *mass-enhanced* 1D Hill-type muscle model to experimentally derived muscle contractions from daily locomotor activities and cycling, and by assessing the effect of muscle inertia on contractile force at human size.

In this present study we use a broad range of movements from slow daily activities to fast, vigorous cycling, which elicit distinct activation patterns across different muscles. By doing so, we can assess both frequency- and size-related effects that would not be evident if only one task or muscle were studied. Therefore, the purpose of this study was to test the effect of including muscle mass in 1D Hill-type muscle models for a variety of human movements and to determine how sensitive these effects are to muscle size. We hypothesized (1) that inertial effects arising from muscle tissue mass would lead to measurable and statistically significant differences in predicted force and power between massless and mass-enhanced 1D Hill-type muscle models for humans undergoing regular activities, and (2) that these effects would increase with muscle size (mass) and acceleration.

## Materials and methods

### Approach to the problem

To determine the inertial effects of muscle mass on muscle force output during *in vivo* human movement, we collected electromyography (EMG) and kinematic data of lower-limb muscles from participants undergoing a range of daily activities (walking and running at a moderate pace, hopping, plus sit-to-stand), and supplemented the data with more data from vigorous cycling that we have reported in a previous study (Dick et al., 2017). By definition, the massless Hill-type model does not include inertial contributions, and thus the geometric scaling factor does not influence the dynamics predicted by this model. Its results are therefore presented solely as a baseline for comparison, with no intrinsic differences expected across scale factors. Any scale-dependent differences observed between models must arise exclusively from inertial effects included in the mass-enhanced model.

A generic musculoskeletal model (Rajagopal et al., 2016) was scaled using motion capture marker data from each participant’s static trial to generate a personalized model. We first scaled the whole-body musculoskeletal model to each participant’s anthropometry, then performed inverse kinematics to estimate muscle–tendon unit (MTU) lengths during the movement tasks. We used the estimated MTU lengths and activation states derived from the EMG data to conduct forward dynamics simulations of muscle force using both massless and mass-enhanced 1D Hill-type muscle models (Fig. 2). Finally, we compared time-varying force, instantaneous power, and net mechanical work outputs from both the massless and mass-enhanced simulations for each trial to isolate (A) the effect of tissue mass, by examining changes across geometric scale factors, and (B) the effect of acceleration, by comparing tasks at different movement frequencies, thereby revealing when inertial forces substantially alter muscle performance.

### Experimental data collection

We collected kinematic, kinetic, and EMG data of lower-limb muscles from twenty participants during a range of daily activity tasks (age (mean±S.D.): ten female and ten male; 33.0±10.7 years old; mass: 66.3±10.1 kg; height: 1.70±0.10 m; details and inclusion and exclusion criteria are described in Table S1). The study protocols were approved by the Institutional Ethics Review Boards at Simon Fraser University. For each participant, 24 LED motion-capture markers were secured bilaterally on the skin over the pelvis and lower extremities, as described in a previous study (Dick et al., 2016). The 3D positions of these markers were sampled at 100 Hz using a dual-head optical motion capture system (Optotrak Certus, NDI, Waterloo, Canada). To record muscle electrical activity, bipolar Ag/AgCl surface EMG electrodes (10 mm diameter, 21 mm spacing; Norotrode; Myotronics, Kent, USA) were positioned over the bellies of the medial gastrocnemius (MG), lateral gastrocnemius (LG), soleus (SOL), rectus femoris (RF), vastus medialis (VM), vastus lateralis (VL) muscles after cleaning with alcohol and shaving the skin. EMG signals were pre-amplified (gain 1000-5000), band-pass filtered (bandwidth 10–500 Hz; Biovision, Wehrheim, Germany), and sampled at 2000 Hz. Ground reaction forces for each foot were recorded at 300 Hz using the dual force plates of an instrumented treadmill (Bertec FP6012-15; Ohio, USA).

Prior to the test session, a static calibration trial was conducted to scale a musculoskeletal model to each participant, based on distances between marker pairs. Participants performed maximal voluntary isometric contractions (MVCs: is the greatest force that a muscle or muscle group can produce during a maximal voluntary effort, usually measured in a standardized isometric test (Takarada & Nozaki 2021)) in the middle of range of motion; the maximum excitation for each muscle recorded from these trials and the test sessions was used for EMG normalization. Participants then continued with a five-minute warm-up by walking on the treadmill to minimize the influence of temperature on the muscle excitations. For the testing trials, participants performed the following four daily activity tasks in order on the treadmill: walking and running at their preferred speed, hopping at a tempo of 110 beats per minute (equivalent to 1.83 s^−1^) timed to a metronome, and sit-to-stand from a chair at the tempo of 15 cycles per minute (equivalent to 0.25 s^−1^). All the data were collected for 30 s for each task. Then, after a 10-minute break, the tasks were repeated in reverse order. One participant was unable to maintain these designated hopping and sit-to-stand frequencies and instead performed hopping at 1.00 s^− 1^ and sit-to-stand at 0.17 s^− 1^.

For the cycling task, kinematic and EMG data were collected from 14 competitive cyclists (7 female and 7 male; 29.9±5.8 years old; mass: 68.8±8.7 kg; height: 1.73.±0.07 m; mean±s.d.) as part of a study of cycling across a range of pedalling conditions, as described in detail elsewhere (Dick et al., 2017). Five pedalling conditions were used in this study, including different crank torque levels of 14, 26, and 44 Nm at a constant cadence of 80 rpm, and cadences of 100 and 140 at a crank torque of 13 Nm. The average crank power outputs for these pedalling conditions were 115 W, 220 W, 370 W, 135 W, and 190 W, respectively.

Intensity envelopes were calculated for the EMG and converted to activation levels (Lee et al. 2011). Activations were subsequently normalized to the activation during maximum voluntary contraction (MVC) trials to yield activations between 0 and 1 for each muscle. The participant-specific LED marker positions were imported to a musculoskeletal simulation environment (OpenSim 4.3: Delp et al., 2007). Participant-specific values of optimal fibre length and tendon slack length were extracted from the musculoskeletal model and balanced to calculate the participant-specific optimal MTU length (equation A1). Using the ‘Inverse Kinematics tool’, we extracted the time-varying MTU length during each task and normalized this value to the participant-specific optimal MTU length.

### Muscle model formulations

A 1D mass-enhanced Hill-type muscle model (Günther et al., 2012; Ross, Nigam, et al., 2018; Ross & Wakeling, 2016) (Fig. 1A) and a traditional, massless Hill-type muscle model (Ross, Nigam, et al., 2018; Ross, Ryan, et al., 2018) (Fig. 1B) were used to simulate muscle contractile behaviours during movement tasks, their formulations are described in the Appendix. In brief, these models consisted of: (1) contractile elements (CE), whose force depended on muscle activation and the active force–length and force–velocity relationships; (2) parallel elastic elements (PEE), producing passive muscle force according to the passive force–length curve; and (3) series elastic element (SEE), representing compliance from the internal and external tendons and aponeuroses, whose force was calculated at each time step from the tendon force–strain length relationship based on the difference between the prescribed MTU (from inverse kinematics) and the current muscle length (Fig. 1C). In the mass-enhanced model, the muscle was evenly split into 16 in-series segments, each of which contained a CE, a PEE, and a point mass that was evenly distributed from the muscle’s total mass (Fig. 1A).

**Fig. 1.**
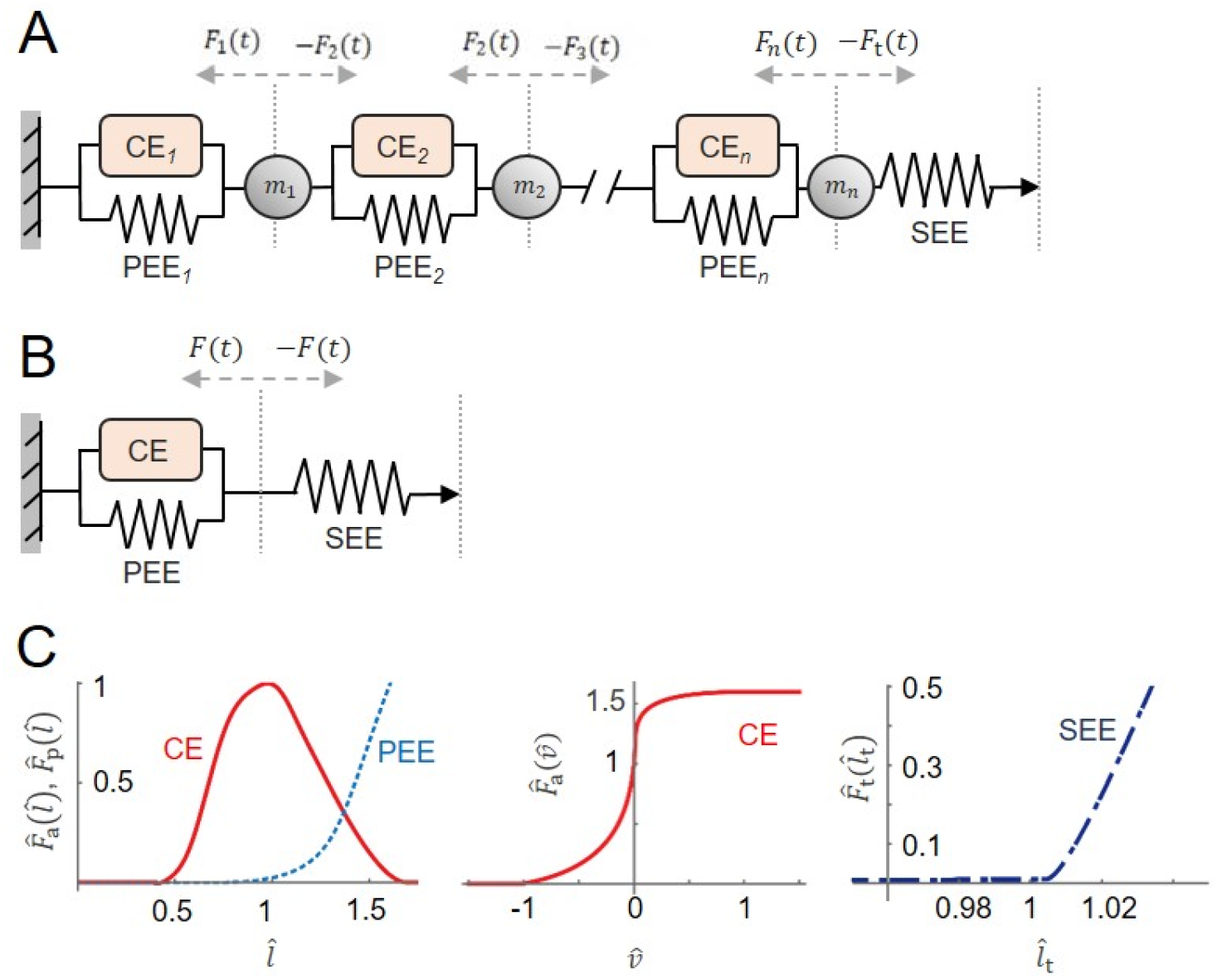
The massless and mass-enhanced Hill-type muscle models used to simulate experimental contraction cycles. The models consisted of an active muscle component (CE, contractile element), a passive muscle component (PEE, parallel elastic element), and a tendon (SEE, serial elastic element) (A) (Wang 2006). The muscle force is the sum of forces from active and passive muscle components, depending on normalized activation level, normalized muscle force-length property (C), and normalized muscle force-velocity property (D). The tendon force depends on tendon force-length property (B). In the mass-enhanced model (E), the muscle is split into segments (=16 in this study) with an optimal segment length evenly distributed from the optimal muscle length, and each segment contains a CE and PEE from the Hill-type model. Between the segments and between the last segment and tendon, there are point masses () divided homogeneously from the muscle mass. Each point mass is accelerated by its adjacent components, either muscle actuators (*F*_*i*_(*t*)) or tendon force (*F*_*t*_(*t*)).*x*_mid_ (*t*), the time-varying position of the muscle-tendon junction; *x*_*i*_(*t*), the position of each point mass; *l*, the muscle length; *l*_*t*_, tendon length; *l*_MTU_, muscle-tendon unit length. The force-length-velocity relationships were piecewise polynomials that were based on Bézier curves (Ross et al., 2018) that had been fitted to experimental data for CE normalized forces (Winters et al., 2011), PEE and force-velocity normalized forces (Roots et al., 2007), and SEE normalized forces (Dick et al., 2016).

**Fig. 2.**
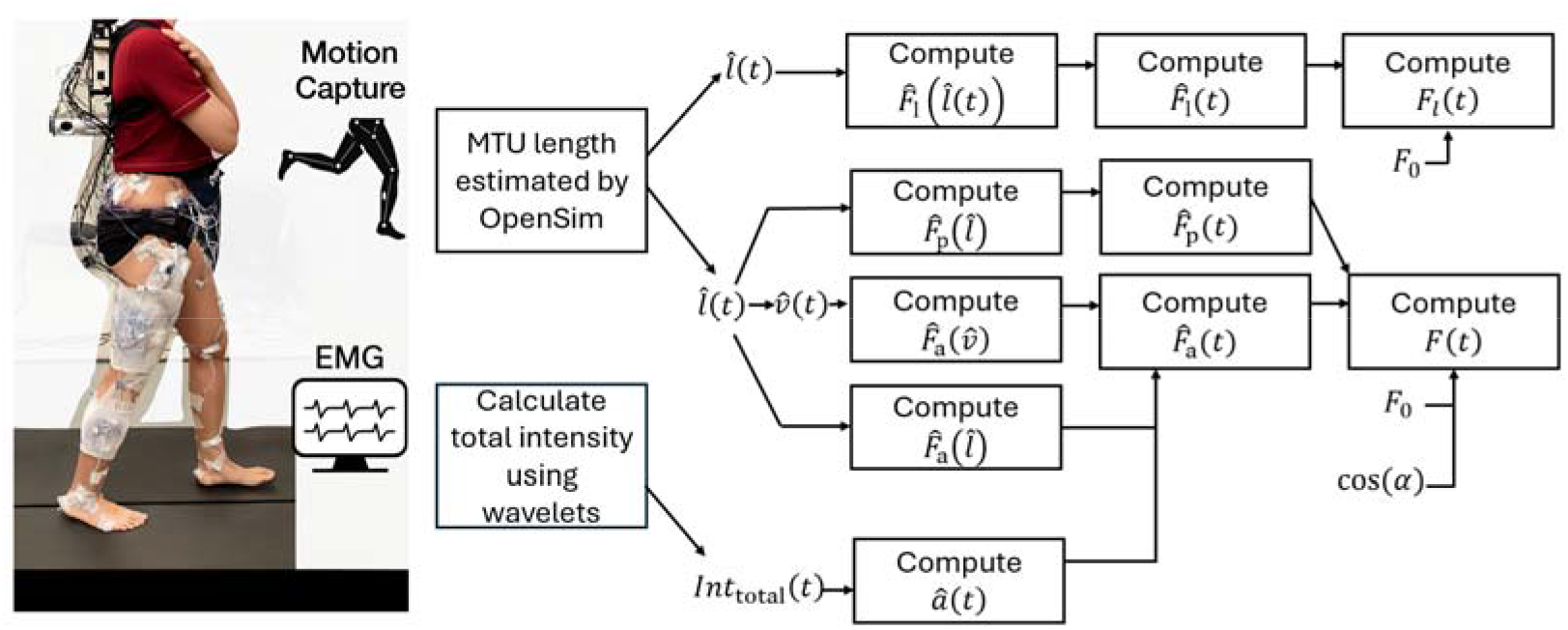
Graphical depiction of data collection and processing. A comprehensive set of kinematic, kinetic, and electromyographic (EMG) data of right lower limb during movement was collected using motion capture system, force plates, and EMG. The time-varying muscle-tendon unit (MTU) length and total intensity of EMG () were estimated using OpenSim and wavelet functions, respectively, and then were used to calculate time-varying normalized muscle length (), normalized tendon length (), and normalized muscle activation level (). These were used as inputs to drive Hill-type muscle models for predicting time-varying muscle force () and tendon force (). : normalized muscle contraction velocity;, normalized tendon force as a function of; and, normalized passive and active muscle forces as a function of, respectively;, normalized active muscle force as a function of; : normalized active muscle force; : normalized passive muscle force; : maximum isometric muscle force; : normalized tendon force.

To simplify geometric scaling and clearly isolate the effects of mass on contractile dynamics, the muscles were assumed to have zero pennation angle; additionally, they had a fixed end, and a free end connected to the SEE. The time-varying MTU lengths, with the activations, were then used to independently drive the massless and mass-enhanced muscle models to predict time-varying muscle forces outside of the full-body simulation framework.

To compare muscles of varying masses while preserving geometric proportions, we defined a dimensionless length scale factor s and applied it so that musculotendon lengths scaled as L⍰ = s·L, physiological cross-sectional areas and maximum isometric forces scaled as A⍰ = s^2^·A and F_0_,⍰ = s^2^·F_0_, and muscle volumes and masses scaled as V⍰ = s^3^·V and m⍰ = s^3^·m. We chose s values of 0.1, 1 and 10 for daily-activity tasks, and 0.1, 0.2, 0.5, 1, 1.6, 2, 3, 4, 5, 6, 7 and 10 for cycling tasks, to span one order of magnitude around the nominal muscle size and thereby isolate the effects of muscle mass on contractile dynamics and inertial forces.

### Comparison of predicted forces

Because the two model types share identical active and passive constitutive laws and differ only by an additive inertial term (Equation A14), they are equivalent in the zero-mass limit (*m*_*i*_ → 0; Equations A7–A8). Consequently, when evaluated with the same MSK-derived kinematics and activations, any increase in RMSD (root mean square difference between the two models’ normalized forces; see Equation A14) reflects the contribution of muscle inertia. Under these conditions, the force predictions and mechanical work output of the mass-enhanced model closely matched those of the traditional (massless) model, with only negligible numerical differences, confirming their mechanical equivalence in the absence of mass. In our analysis, we computed both the RMSD and the coefficient of determination (r^2^) between the forces predicted by the massless and mass-enhanced models across trials for various tasks, including five cycles each of walking, running, and hopping, one cycle for sit-to-stand, and three cycles for cycling.

For cycling specifically, we evaluated the net muscle-volume-specific work output 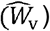 and mean muscle-volume-specific power output 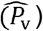 per cycle and contrasted these between the mass-enhanced and massless model predictions over varying muscle sizes. Here, both VL and soleus muscles were examined due to their relatively large volume proportions within the quadriceps and triceps surae muscle groups, respectively.

In order to assess the relative contribution of inertia and activation, we performed a comparison of how the predicted normalized forces interact with muscle activation (Equation A7), influencing observed errors. Prior work showed that the presence of inactive fibres exacerbates the mass effect and error in massless models during submaximal contractions (Ross and Wakeling. 2016). By comparing the equations that describe tendon force predictions from the massless (Equation A11) and mass-enhanced models (Equation A14), we determined that the RMSD differences (Equation A16) arise from two key components: one involving activation and the normalized kinematics (length and velocity) and the other involving the inertial contributions (scaled with muscle mass and acceleration). Although activation influences both, the impact of mass and acceleration appears dominant.

### Statistics

To test the influence of muscle mass on force predictions, we performed separate statistical analyses (IBM SPSS Statistics, Version 27) for (a) activities of daily living and (b) cycling tasks. Because the variance of the RMSD increased with scale factor, a log-transformation was applied to the RMSD values to meet assumptions of homogeneity. We used general linear model ANOVAs test the effect of muscle mass on the log-transformed RMSD force and power, using length scale, movement condition, and muscle as fixed factors, and including the participant as a random factor. Statistical significance was set at p<0.05. Post-hoc pairwise comparisons were performed when significant main effects or interactions were found (Meier, 2006).

## Results

A total of 530 movement cycles were analyzed from each of the 20 participants who performed daily activity tasks (100 cycles each from walking, running, and hopping, and 20 cycles from sit-to-stand, also, a total of 210 cycles were analyzed for cycling tasks). The walking and running speeds were 0.73±0.20 m s^−1^ and 1.51±0.30 m s^−1^ (mean ± S.D.), respectively, while their corresponding cadences (stride frequency) were 0.79±0.10 s^−1^ and 1.30±0.10 s^−1^ (mean ± S.D.), respectively. The cadence (movement frequency) for hopping and sit-to-stand tasks were initially set at 1.83 s^−1^ and 0.25 s^−1^, respectively, but could be adjusted according to participants’ preference. The average hopping and sit-to-stand cadences were 1.79±0.19 s^−1^ and 0.25±0.02 s^−1^. For the cycling tasks, the cadences were set at 1.33 s^−1^, 1.67 s^−1^, and 2.33 s^−1^, for 80, 100, and 140 rpm, respectively. Higher cadence (high frequency) movement leads to a higher overall muscle average acceleration in general (Fig. 3).

**Fig. 3.**
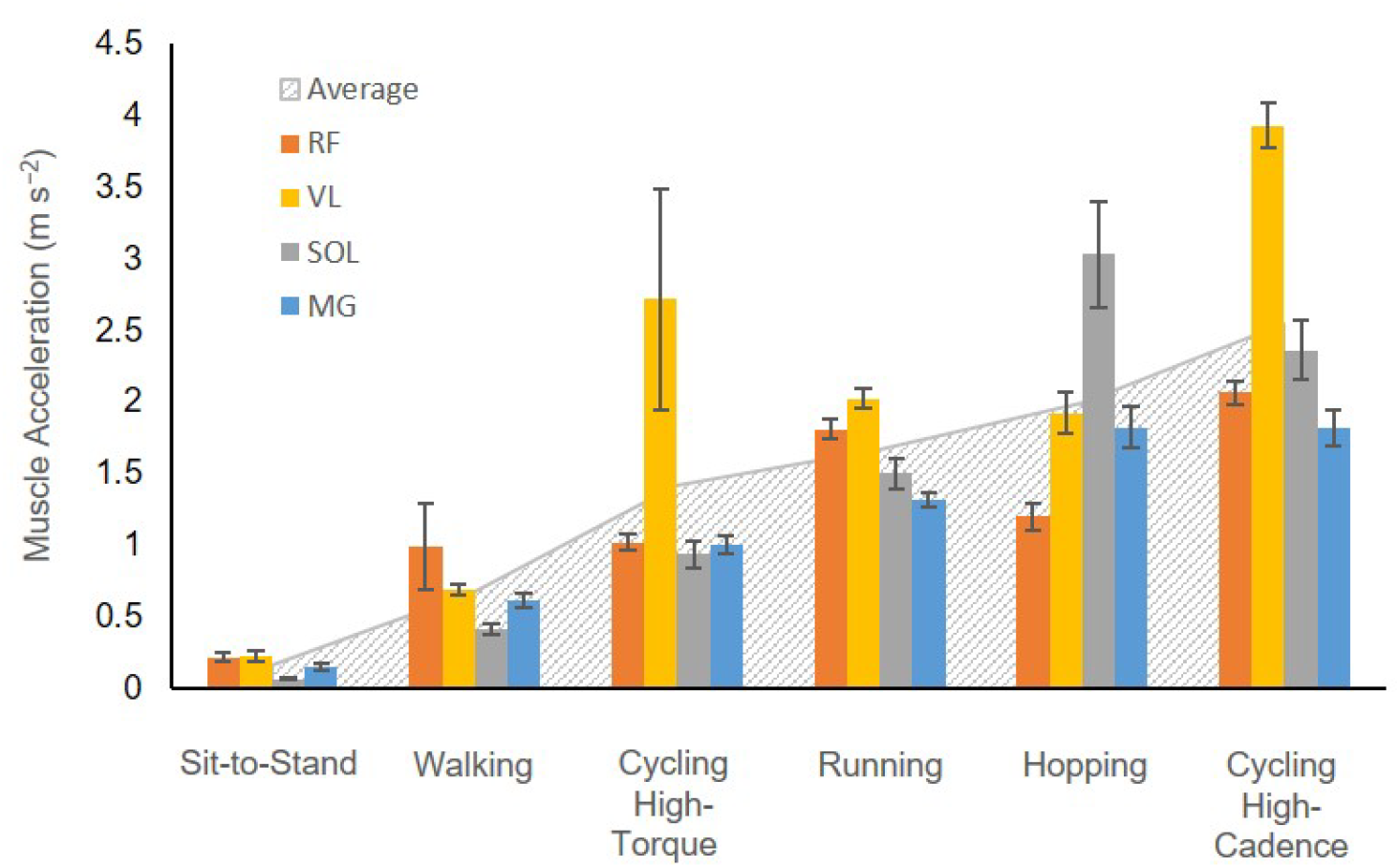
Muscle accelerations in the tested muscles across tasks at scale 1. Solid bars show the mean ± s.e.m. of root-mean-squared muscle accelerations per cycle, while the boundary of grey dotted background represents the muscle acceleration averaged across muscles in each task. Tasks are ranked by the average muscle acceleration, which correlates with movement cadence. This indicates that movements with higher cadence (or frequency) result in greater average muscle acceleration overall. High-torque cycling refers to conditions of 44 Nm and 80 rpm, while high-cadence cycling refers to 140 rpm and 13 Nm.

The predicted normalized time-varying forces, 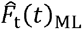 for the massless model, showed no variation beyond numerical error despite changes in scale (0.1, 1, and 10). These values also closely resembled the 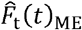 predicted by the mass-enhanced model at scale 0.1 and 1, leading to substantial overlap between the predicted 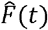 from both models (Fig. 4B) at these sizes. By contrast, a notable difference emerged at scale 10 (Fig. 4B, bottom row), where the predicted 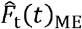 from the mass-enhanced model showed fluctuations that were periodically higher or lower than that of the massless model.

**Fig. 4.**
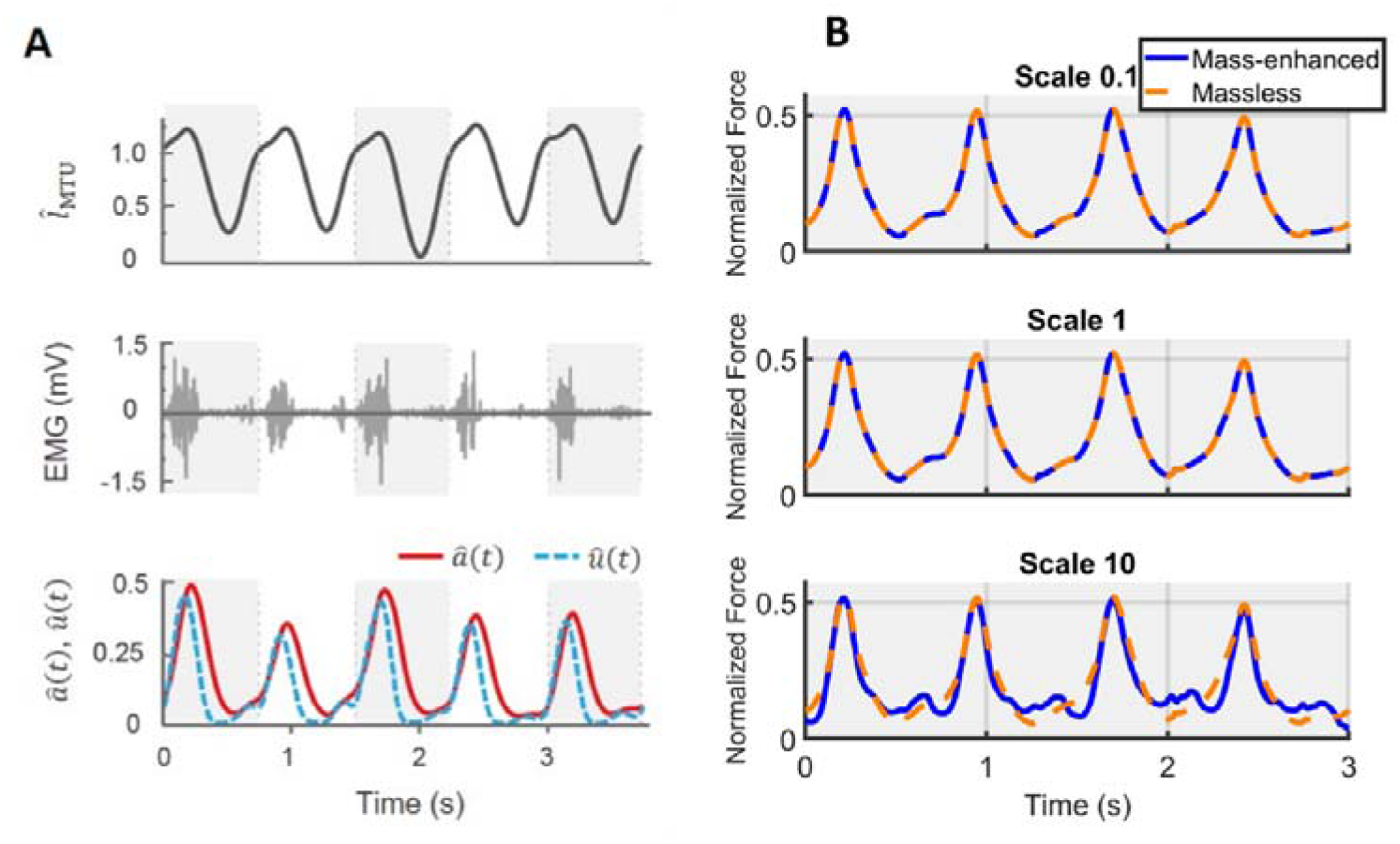
Sample raw traces and predicted muscle forces using massless and mass-enhanced muscle models at different scales. Muscle-tendon unit (MTU) length normalized to its optimal length over time is shown in the top row of (A). Each grey or white block represents one movement cycle. The middle row shows raw electromyographic (EMG) data over time. The EMG data are converted to muscle excitation normalized to the maximum excitation (û(*t*)) (blue, dashed) and muscle activation normalized to the maximum activation (â(*t*)) (red, solid) over time, which are shown in the bottom row. Using time-varying normalized MTU length and normalized activation level inputs, normalized muscle force 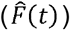 over time is computed with massless or mass-enhanced model. (B) depicts predicted time-varying muscle forces from both models across three scales. The predicted forces from both models overlap at scales 0.1 and 1, but at scale 10, a noticeable difference in predicted forces emerges between the models. The data are taken from right medial gastrocnemius MG muscle of one representative participant during the running task.

The ANOVA tests for the log-transformed RMSD showed a significant effect of scale and movement type on the model predictions (Fig. 5). There was a significant effect of the participant factor for the activities of daily living, but not for the cycling conditions. Larger scales consistently exhibited higher log-transformed RMSD forces, regardless of the muscles or movement tasks involved. When further looking into the relations between scale factor and RMSD forces for the high-cadence cycling data, the log-transformed RMSD in 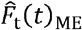 showed a monotonic increase with the log-transformed scale factor across all tested muscles (Fig. 6A). Additionally, the log-transformed RMSD in 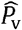 also showed an increase with the scale factor (Fig. 6B). When disaggregating the data based on sex, there was no significant effect of sex on the log-transformed RMSD in 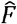 (Fig. 1).

**Fig. 5.**
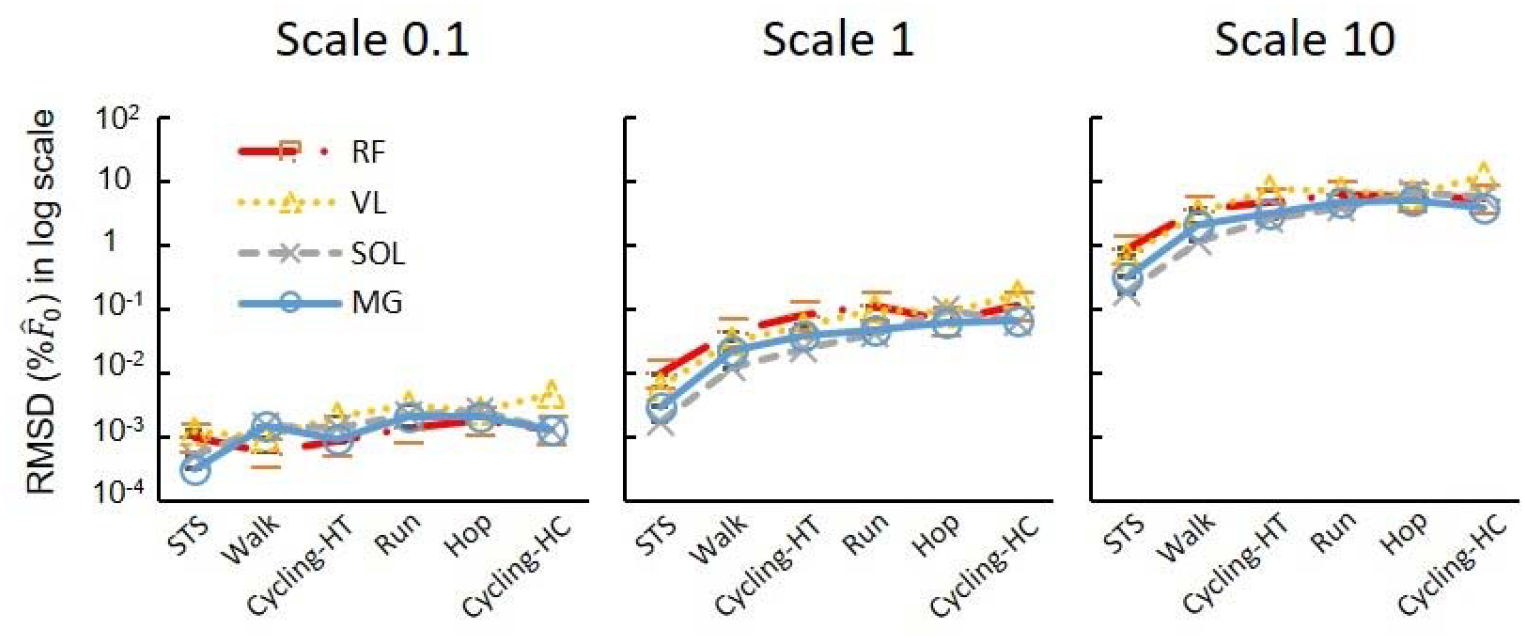
The effects of scaled muscle mass (size) on the difference between model-predicted forces. Each plotted point denotes the average of log-transformed RMSD in 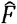 across participants for a particular muscle engaged in a specific task and scale. When comparing across scales, larger scales consistently exhibited higher log-transformed RMSD in 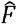, regardless of the muscles or movement tasks involved. High-torque cycling refers to conditions of 44 Nm and 80 rpm, while high-cadence cycling refers to 140 rpm and 13 Nm.

**Fig. 6.**
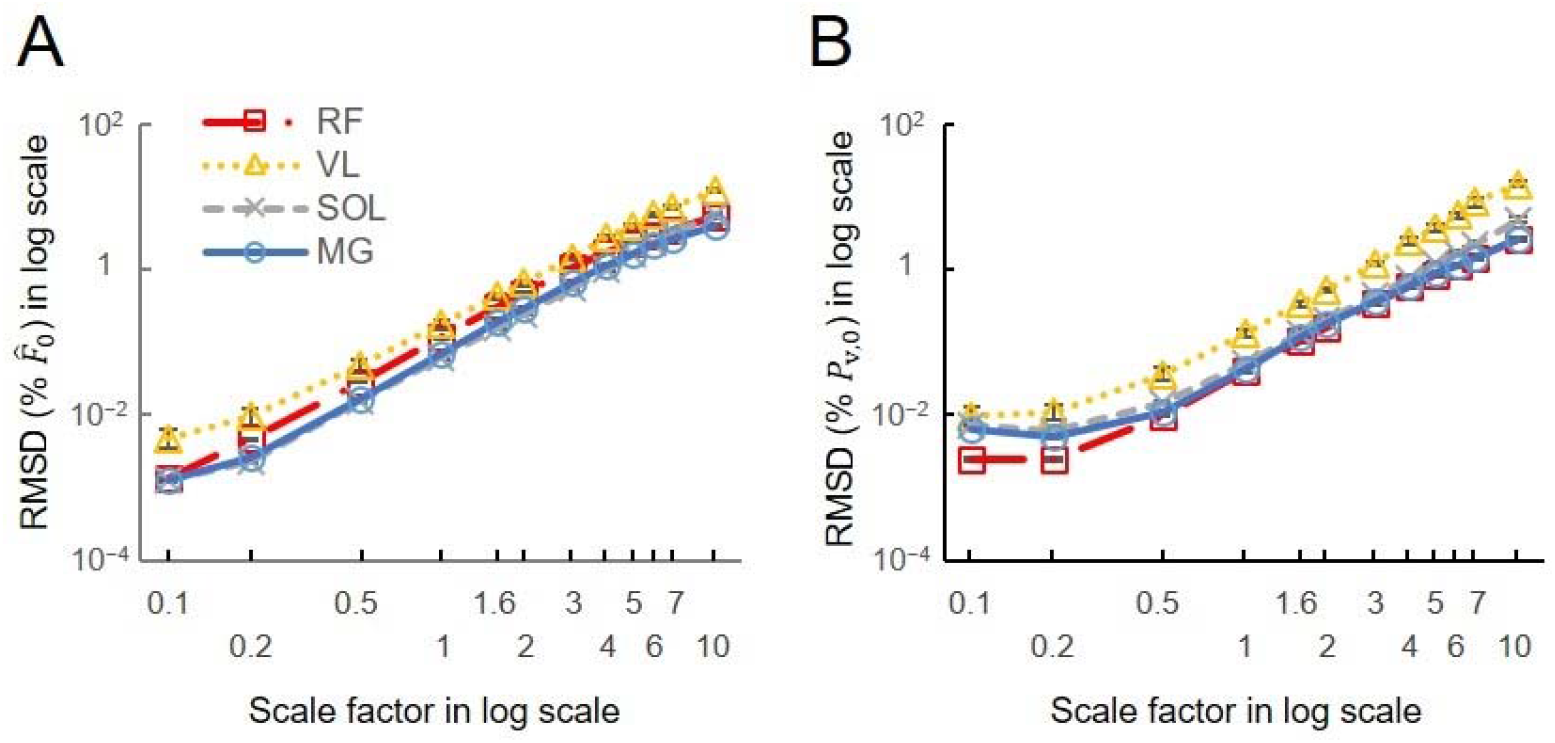
The effects of scaled muscle mass (size) on the difference between model-predicted normalized forces and volume-specific power. (A) The log-transformed RMSD in 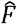 shows a nearly linear increase with the scale factor across all tested muscles (rectus femoris RF, vastus lateralis VL, soleus SOL, medial gastrocnemius MG), indicating that 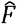 predicted by mass-enhanced model deviate more distinctly from the 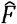 predicted by massless model at scale 1 when scaled muscle size becomes larger. (B) The log-transformed RMSD in *P*_v,o_ also shows a increase with the scale factor. The representative data was taken from muscles in high cadence (140 rpm)-low load (13 Nm) cycling condition averaged across participants.

The RMSD in 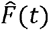 and standard error values were consistently higher in high-cadence movement tasks (running, hopping, and both cycling tasks) (Fig. 5), compared to lower cadence tasks (walking or sit-to-stand tasks) for any given muscle at scale 1 (*P*<0.05).

Variations between model predictions increased with cycling cadence (significant effect of cadence: P<0.05) but were not significantly influenced by the crank load (Fig. 7). RMSD in 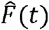 increased with cycling cadence when the torque was set at 13-14 Nm. Averaged across scales, the RMSD in 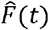 for VL muscle were 1.60±0.11, 2.47±0.20, 3.33±0.21 % 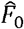, for 80, 100, 140 r.p.m., respectively, with significant differences found among these cadences (*P*<0.05) when the crank load was 13-14 Nm. In contrast, when the crank load increased from 14-44 Nm (at a fixed cadence of 80 rpm) there was no significant effect of load on the RMSD_ME_ for 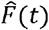.

**Fig. 7.**
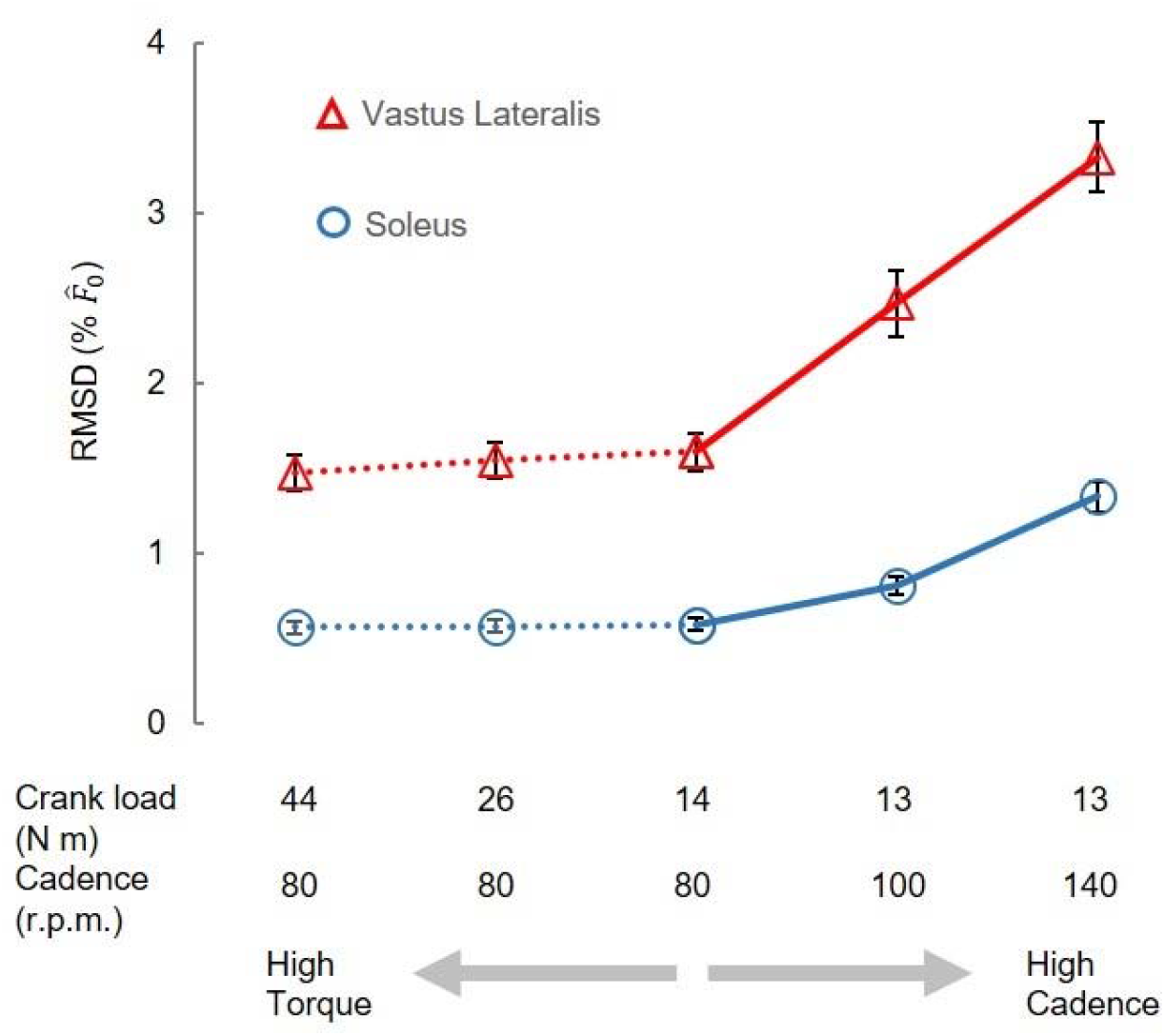
Comparison of mass effects among cycling conditions. The RMSD values in vastus lateralis (VL) and soleus (Sol) muscle, when averaged across scales, increases significantly with cycling cadences. The RMSD values in VL and Sol muscle are not significantly different among various crank loads.

For the cycling tasks, the volume-specific net, positive, and negative muscle work outputs per cycle predicted by the massless model remained relatively constant for both the VL (5.49, 10.50, −5.01 kJ m^−3^, respectively) and the SOL (2.37, 6.47, −4.10 kJ m^−3^, respectively) muscles, despite the increasing scale factor (Fig. 8). In contrast, the volume-specific net, positive, and negative muscle work outputs per cycle predicted by the mass-enhanced model exhibited variability across scales. At smaller scales (scale factor 3 or less), these outputs mirrored those of the massless model, remaining relatively constant. However, beyond a scale factor of 4, there was a noticeable trend: the volume-specific positive work tended to decrease, while there was an increase in the magnitude of the volume-specific negative work. As a result, the volume-specific net work of the muscle tended to become smaller as muscle mass increased. Including muscle mass in the simulations reduced the net mechanical work output per cycle, compared to the massless model, due to the inertial resistance of the muscle segments. When normalized to muscle volume, this volume-specific net work decreased more steeply with increasing cadence than with decreasing crank load for both the vastus lateralis and soleus muscles (Fig. 9). Additionally, the difference in volume-specific net work between the mass-enhanced and massless muscle models became more pronounced with higher cadence and larger muscle scale factors, highlighting the growing impact of internal inertial forces under faster and larger muscle conditions.

**Fig. 8.**
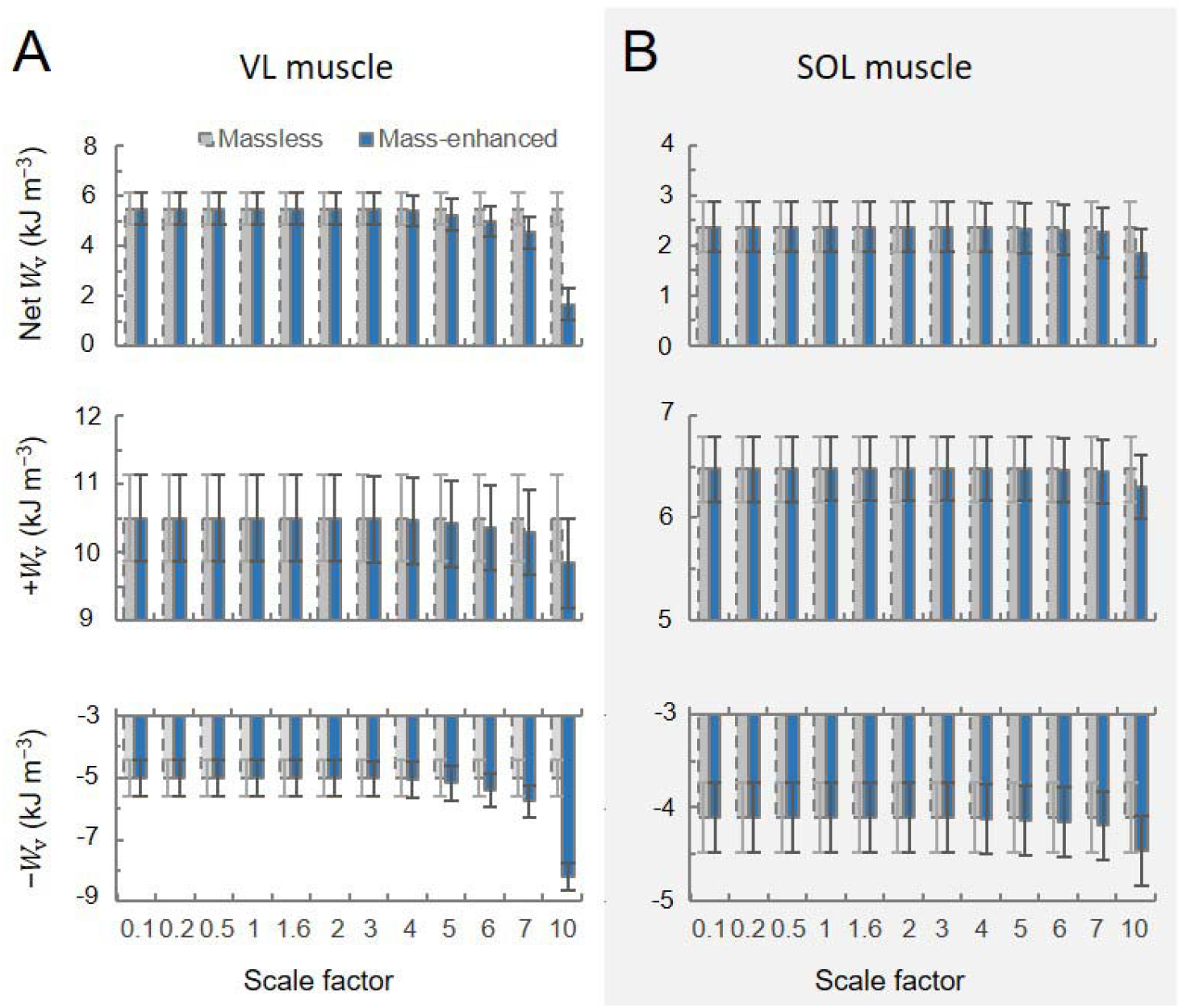
Volume-specific muscle work output per cycle (*W*_v_) across scales during high torque (44 Nm)-low cadence (80 rpm) cycling condition. The top, middle, and bottom rows represent net, positive, negative muscle work output per cycle across scale factors, respectively, for the vastus lateralis (VL) (A) and the soleus (SOL) (B) muscles. Work outputs were predicted using both massless (grey bar) and mass-enhanced (dark blue bar) muscle models.

**Fig. 9.**
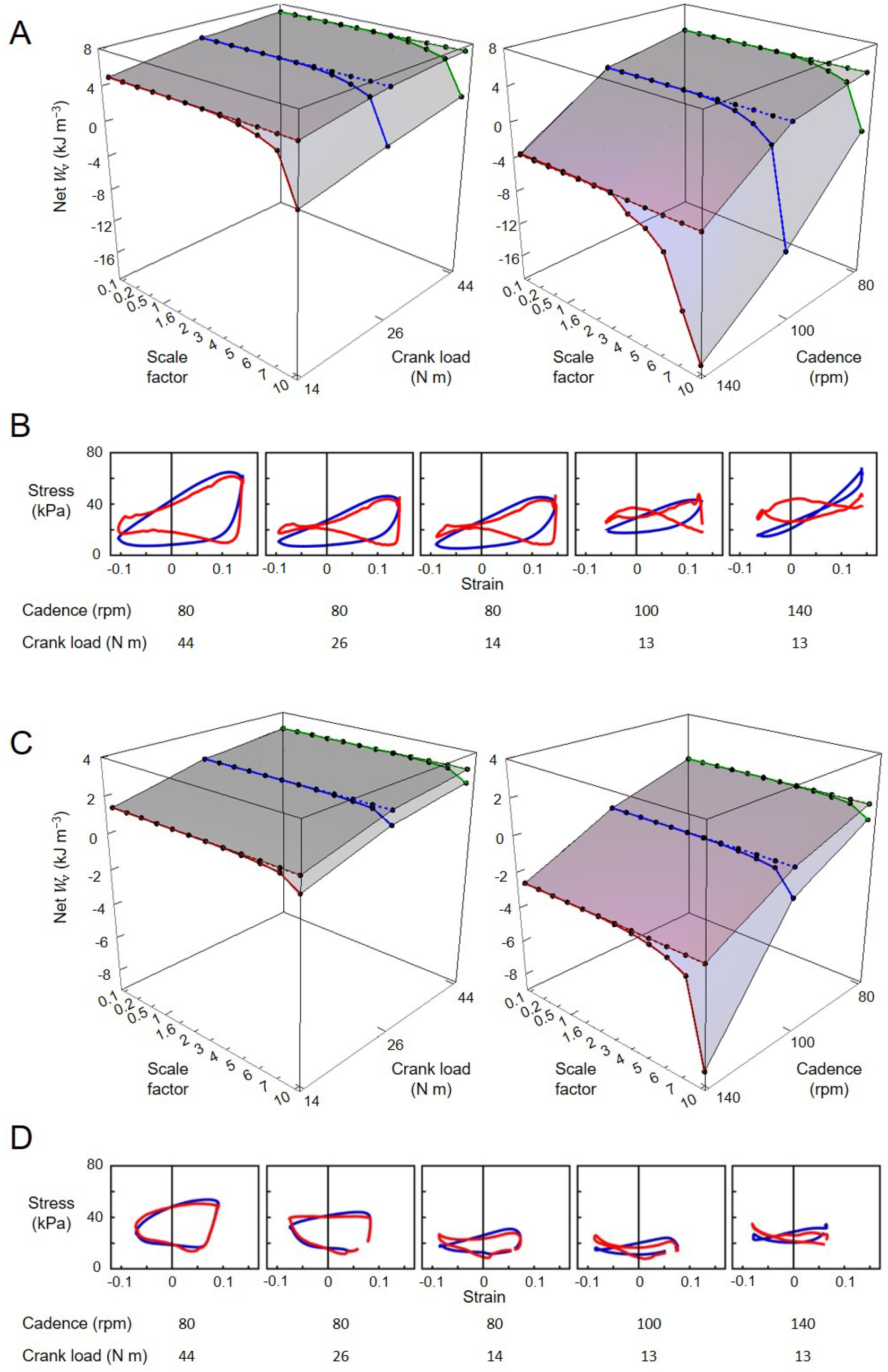
Volume-specific muscle net work output per cycle (*W*_v_) across scales and cycling conditions. (A, C) Each point represents *W*_v_ at a specific scale factor and cycling condition. Points connected by a dotted line indicate net *W*_v_ calculated using the massless muscle model, while those connected by a solid line represent net *W*_v_ calculated using the mass-enhanced model. Graphs B and D are volume-specific work loops across cycling conditions. Blue loops are derived from the massless model while red loops from the mass-enhanced model. Graphs A and B show data from the vastus lateralis muscle, while C and D are from the soleus muscle.

## Discussion

The purpose of this study was to examine the impact of modelling muscle mass and inertial effects on muscle force output during *in vivo* human movements. We tested whether the inclusion of distributed muscle mass in mass-enhanced 1D Hill-type muscle models would affect the predictions of muscle force during human movements and if scaling muscle size would alter this effect. Overall, our results showed that while a massless Hill-type muscle model predicted the same normalized forces across the range of scales tested, a mass-enhanced Hill-type muscle model showed an increasing effect of mass as the muscle size increased and as the cadence and thus acceleration increased. These results coincide with previous studies examining the impact of muscle tissue inertia on the mechanical output of isolated muscle *in situ* in rats (Ross et al., 2020) and *in silico* using 1D and 3D muscle models (Günther et al., 2012; Meier and Blickhan, 2000; Ross et al., 2021; Ross, Ryan, et al., 2018; Ross and Wakeling, 2016, 2021). However, the results show that the magnitude of the effect of muscle mass on the forces predicted by 1D mass-enhanced Hill-type muscle models is small (< 1 %) for human-sized muscles during human movements, particularly the slower movements of daily activities. Given the additional complexity of computing the mass dynamics within a muscle, benefits of including mass in the dynamics of predicting human muscle force seem elusive in this context.

The significant effect of the participant factor on the RMSD force observed during activities of daily living is likely caused by the variation in locomotion patterns between the participants. The walking and running tasks were completed at the participant’s preferred speeds and the hopping and sit-to-stand tasks were completed at different cadences between participants. These differences would have led to differences in the accelerations experienced by the muscles and thus participant-specific differences in the effect of mass. On the other hand, the cycling trials were much more constrained, being at specified cadences and crank torques, and with the lower limb segment trajectories constrained by the path of the pedals. Indeed, there was no significant effect of the participant factor on the RMSD forces for the cycling conditions.

Muscle mass affects contraction dynamics because there is an inertial cost to accelerating that mass, and the energy required to accelerate the muscle mass takes away from the energy available to do external mechanical work. Furthermore, as size increases, the muscle mass (scaling with the cube of the length-scale factor) would increase more than the force available to accelerate it (scaling with the square of the length-scale). Thus theory predicts greater mass effects at larger sizes (where the mass is higher), faster movements (where the accelerations may be greater), and lower levels of activation (where there would be less force). In this study, we compared the effects of tissue inertia during five movements that cover a range of cadences and consequently muscle accelerations: sit-to-standing, walking, running, hopping and cycling. The results reflect the model’s physical assumptions, namely, that increased accelerations of muscle mass require greater inertial forces from the mass-enhanced muscle models. The task-level application in this study extends previous findings from abstracted muscle models and helps characterise when and where mass effects become most functionally relevant in human movement.

In the gait conditions we examined, movement speed is typically coupled with muscle activations such that faster tasks, like running, result in higher muscle activations (Biewener & Patek, 2018). To uncouple speed and activation more fully we probed the cycling data more deeply, because during cycling, unlike walking and running, speed and load can be varied independently. The increased mass effects at higher cadences become more noticeable as scaled muscle size increases, especially beyond human-sized muscles (Fig. 6A). Our results showed that when predicting forces of a muscle 6^3^ times larger than its original mass, mass effects could lead to errors exceeding 5 % at a pedalling cadence of 140 r.p.m., while remaining below 5 % at lower cadences (Fig. 6A). However, for smaller muscle sizes, mass effects appeared less sensitive to pedalling cadence. Equation A16, which elucidates the variables influencing RMSD calculations, highlights that the scale factor and pedalling cadence, directly linked to mass and acceleration magnitudes, primarily determine these differences. In contrast, crank load, which impacts activation levels to a greater extent than cadence (Dick, Arnold, and Wakeling 2016), showed no significant effect on the differences between the models (Fig. 7).

The mass effects identified in this study likely underestimate the actual mass effect occurring *in vivo*, given that our models were constrained to a single dimension and thus omitted multi-directional deformation and acceleration of muscle tissue. The mass effects that we predicted were small for the human-sized models at scale 1, and for the relatively slow activities of daily living that we examined in this study. The largest mass effect we observed amounted to 7.23 % 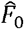 (with a *r*^2^ of 0.68), and corresponds to the RMSD (and coefficient of determination) in predicted force between the mass-enhanced and massless models at a larger than human-size scale 10 for the vastus lateralis during running. In the 1D models in this study, the distributed tissue mass was constrained to only allow for movement along the muscle’s length. However, in real muscle the tissue also deforms and bulges in width and thickness such that its mass accelerates in all three dimensions. In previous work using a 3D muscle model to examine mass effects in isolated muscle, we found up to 34.4% reduction in mechanical work per sinusoidal contraction cycle at a scale of 3.5 times greater than the human medial gastrocnemius (Ross et al., 2021). In addition to the dimensionality of the muscle model, there are other factors that could explain discrepancies in mass effects between studies. In Ross et al. (2021), the maximum muscle size was smaller than that in the current study (scale of 3.5 compared to 10), and all the simulated 3D muscles were pennate which could decrease the effect of mass. The mean root-mean-square accelerations in the Ross et al. study may have been higher (ranged from 2.4 m s^-2^ for strain amplitude of 2.5% to 7.1 m s^-2^ for amplitude of 7.5%) than in this study (Fig. 7) which could increase the mass effects. However, it is likely that these factors would have a relatively small effect on predicted force output compared to dimensionality differences.

We report here the effect of muscle mass on the muscle force, due to the test being kinematically constrained with the trajectories of the MTU length varying only with model size, and not with model type. Previously we have shown that smaller mass effects occur when mass-enhanced muscle models contract against damped harmonic oscillators (where both displacement and force can be changed) as compared to kinematically constrained sinusoidal contractions (Ross & Wakeling, 2021). However, because harmonic oscillators can only store and return energy and cannot input energy into the system like a motor, the restoring forces were too small to substantially lengthen the muscle during the relaxation phase of the contraction cycle. As a result, the muscle strains were either much smaller or at shorter lengths for the harmonic oscillator compared to the sinusoidal simulations; this likely contributed to the smaller mass effects for the unconstrained contractions against the harmonic oscillator. Thus, it is possible that greater mass effects would occur than those we found in this study in predictive simulations of movement when mass-enhanced models have the ability to affect both the kinematics and kinetics of the movement and when larger restoring forces are provided by antagonist muscles.

The inclusion of a tendon in mass-enhanced models has been shown to mitigate the decline in mechanical work output per cycle associated with increased mass (Ross and Wakeling, 2021). Each modeled MTU had an optimal tendon stiffness that maximises work output, as previously identified by (Ross and Wakeling 2021), and this optimal stiffness is influenced by scaled muscle sizes and the timing of muscle excitation in relation to the start of MTU shortening (Ettema, 2001; Lichtwark and Barclay, 2010; Lichtwark and Wilson, 2005; Sawicki et al., 2015). We utilised a standardized, non-dimensional series elastic element (SEE) force–strain relationship across all muscles, as commonly adopted in musculoskeletal simulations, resulting in muscle-specific absolute tendon stiffness values dependent on each muscle’s tendon slack length and maximal isometric force (Delp et al., 2007). We acknowledge that muscle-specific tendon stiffness may affect the work done by the muscles and the absolute mass effect, however, the simplifying assumption of uniform tendon stiffness should not affect the main findings of differences in predicted forces between mass-enhanced and massless models, and an increasing influence of muscle mass with the size of the model.

Despite the valuable insights gained from our 1D modelling approach, several limitations should be acknowledged. First, our results rely exclusively on simulation predictions without direct experimental validation of muscle forces or mechanical work output. Second, using inverse kinematics and prescribed musculotendon-unit (MTU) lengths precludes feedback from muscle dynamics into joint motion, potentially oversimplifying musculoskeletal interactions. Third, model simplifications, including zero pennation angle, fixed tendon properties, and uniform geometric scaling may restrict generalization to real muscle conditions. In vivo muscle is anisotropic: fibers insert at nonzero pennation angles that vary with length and activation, and aponeuroses form compliant, load-bearing sheets with spatially nonuniform strain fields. Dynamic pennation and architectural gearing can alter fiber shortening velocity relative to musculotendon excursion, thereby changing the balance between active force production, power, and series-tissue loading; neglecting these effects may bias line-of-axis force transmission (potentially overestimating it when pennation is ignored) and modify the apparent magnitude of mass-related differences. Moreover, aponeuroses do not deform uniformly: their strain distributions depend on contraction mode (shortening vs lengthening), activation level, and transverse pre-tension, leading to heterogeneous stress and shear that our simplified series elastic representation does not capture (Raiteri 2018, Böl et al. 2015). These architectural features can change fiber velocities, internal work, and energy storage/dissipation depending on task and geometry, but it is not easily predictable whether these effects will be increases or decreases.

Future research should address these inherent limitations by extending simulations into three dimensions. Developing 3D muscle models would allow us to capture multi-dimensional deformations and complex inertial effects, such as tissue bulging and nonlinear mass distributions, that occur *in vivo* (e.g., via 3D or finite element models) to quantify how architectural anisotropy interacts with tissue mass and movement speed to shape contractile performance. Moreover, rather than focusing on a limited set of muscles, future studies should aim to simulate the entire human lower limb musculoskeletal system. Such comprehensive models could incorporate the intricate interplay of muscle activation patterns, joint kinetics, and antagonistic interactions that influence overall movement performance and mechanical work output. These advanced simulations would not only refine our understanding of muscle dynamics under various conditions but also enhance the predictive capability of musculoskeletal models for clinical applications, athletic performance assessment, and rehabilitation strategies.

## Conclusions

This study marks the first to use a mass-enhanced muscle model for simulating human muscle contractions during a range of locomotor activities. When we scaled muscles to different sizes, we discovered that mass effects, observed through the RMSD between predicted forces in a 1D mass-enhanced muscle model and a massless model, were minor for smaller muscles but became more pronounced as the muscles exceeded human size. Our analysis quantifies the directional component of mass-related effects in a 1D setting. Allowing three-dimensional tissue motion and pennation could reduce accelerations required along that axis (diminishing directional inertial penalties) yet introduce transverse and shear dynamics that alter fibre velocity, power, and internal work. Consequently, the net in vivo impact of tissue mass and inertia is task- and architecture-dependent rather than uniformly larger or smaller than our estimates. Within our simulations, predicted mass-related differences were larger for running, hopping, and cycling than for walking and sit-to-stand across the muscles tested. These model-based patterns are consistent with the higher movement speeds/accelerations in those tasks, but should be interpreted as quantitative predictions of this modelling framework rather than direct evidence of in-vivo effects; empirical validation will be needed to confirm their magnitude in practice. These findings suggest that while mass effects can significantly influence muscle performance during daily human movement, its impact on forces predicted by 1D muscle models may be minimal in human-sized or smaller muscles during slower activities. However, inertial effects become more apparent at faster cadences and larger sizes.

## Acknowledgements

The authors would like to express their gratitude to Sidney Morrison, Hannah Wood and Nicole Conquergood for their assistance with the experiment. The authors thank all the volunteers for participating in this study.

## Appendices

### Optimal lengths for muscle model

To determine the participant-specific optimal MTU length (*l*_O,MTU_), we used values for optimal fibre length (*l*_f,O,OpenSim_), slack tendon length (*l*_t,slack,OpenSim_), and pennation angle (*l*_OpenSim_), which were extracted from musculoskeletal models (OpenSim 4.3) scaled to each participant, and calculated as follows:

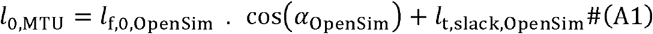

To determine the participant-specific optimal muscle and optimal tendon lengths, we first calculated an estimated optimal muscle length (*l*_O,est_) and an estimated optimal tendon length (*l*_t,O,est_) as follows:

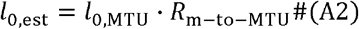

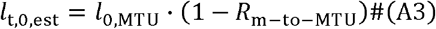

where *R*_m-to-MTU_ is the muscle-to-MTU-length ratio for particular muscle (Kovács et al., 2020; O’Brien et al., 2010; van der Made et al., 2015). Then, we assumed that the actual optimal muscle length (*l*_O_) and optimal tendon length (*l*_t,O_) were “off” by a factor *e*, that is:

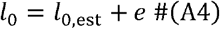

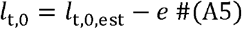

For each participant and muscle, the error term *e* was determined by balancing muscle and tendon forces under zero activation:

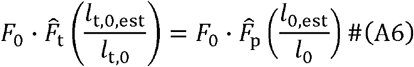

where 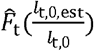 is the normalized tendon force as a function of normalized tendon length and 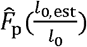 is the normalized passive muscle force as a function of normalized muscle length; *F*_O_ is the maximum isometric muscle force. The maximum isometric muscle force was calculated as the physiological cross-sectional area multiplied by the maximum isometric muscle stress (*σ*_O_= 225 kPa; estimated from literature of Medler, 2002), where the physiological cross-sectional area equals individual muscle volume (Handsfield et al., 2014) divided by the optimal muscle length.

### Massless and mass-enhanced Hill-type muscle models

Let *l*(*t*) and *l*_*t*_(*t*) denote the (unnormalized) muscle belly and tendon lengths, respectively. Muscle force can be calculated by solving the following equations for a massless Hill-type muscle:

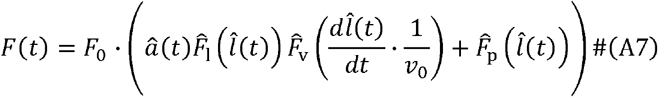

Where 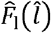 is the active force–length relationship (unitless), 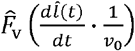 is the active force–velocity relationship (unitless), and 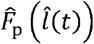 is the passive force–length relationship (unitless).

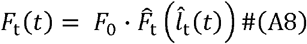

where 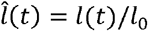 and 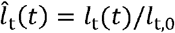 are the normalized muscle belly and tendon lengths, respectively, and *F*_O_ is the maximum isometric muscle force.

The mass-enhanced Hill-type muscle model incorporated muscle mass using 16 point masses distributed along the muscle’s length (Ross et al., 2018a). These masses were connected by Hill-type actuators that generated force to accelerate formulation as the massless Hill-type model, each one with a segment length *l*_*i*_(*t*). them. The segments between point masses were modelled using the same The last point mass at the free end of the muscle was connected to a massless tendon. Each point mass was accelerated by the resultant of its adjacent forces (*F*_*i* + 1_ (*t*), *F*_*i*_ (*t*), or *F*_*t*_(*t*)), which changed the position of each point mass (*x*_*i*_(*t*)). For each point mass, the resultant of the forces from the neighboring segments (*F*_*i* + 1_ (*t*) and *F*_*i*_(*t*)) equaled to the product of mass (*m*_*i*_) and its acceleration caused by that resultant force. For the last point mass (*m*_*n*_), the resultant force acting on it came from the force of the *n*^th^ segment (*F*_*n*_ (*t*)) and the tendon force (*F*_t_ (*t*)):

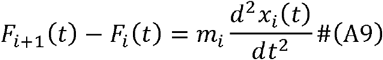

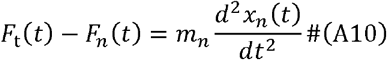

where *i* is an integer ranging from 1 to 15, denoting the point mass and segment number, while *n* is 16, representing the last point mass and segment.

### Relative effect of activation, mass and acceleration on predicted forces

The predicted normalized muscle forces interact with the level of muscle activation (Equation A7), potentially influencing the observed errors. A previous study has demonstrated an increase in errors in traditional massless Hill-type models when estimating in situ cat soleus muscle forces during submaximal contractions compared to maximal contractions (Millard et al. 2013). One explanation is that inactive fibres in submaximal contractions contribute to internal load instead of producing force, thus amplifying the mass effect and increasing the errors in massless models. To understand how muscle activation affects RMSD 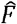 values, we further calculated the difference in 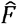 predicted by mass-enhanced and massless models using the aforementioned equations. We assumed that muscle forces were entirely transmitted to the tendon, equating muscle forces to tendon forces in both models. By utilizing Equations A7–A8, the tendon forces of the massless model (*F*_*t*_ (*t*)_ML_) were predicted as follows:

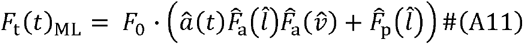

To calculate tendon forces (*F*_t_ (t)_ME_) in the mass-enhanced model, we combined Equations A9 and A10 as follows:

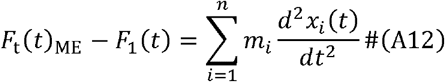

Subsequently, by substituting *F*_1_ (t) (the first muscle segment force) with Equation A7, the tendon forces were calculated as follows:

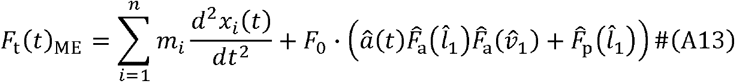

Where 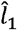 and 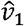 were the normalized length and velocity of the first muscle segment:

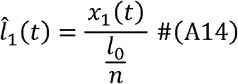

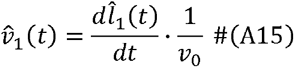

The values of RMSD between model-predicted normalized muscle forces were therefore determined by the differences between Equations A11 and A14 as follows:

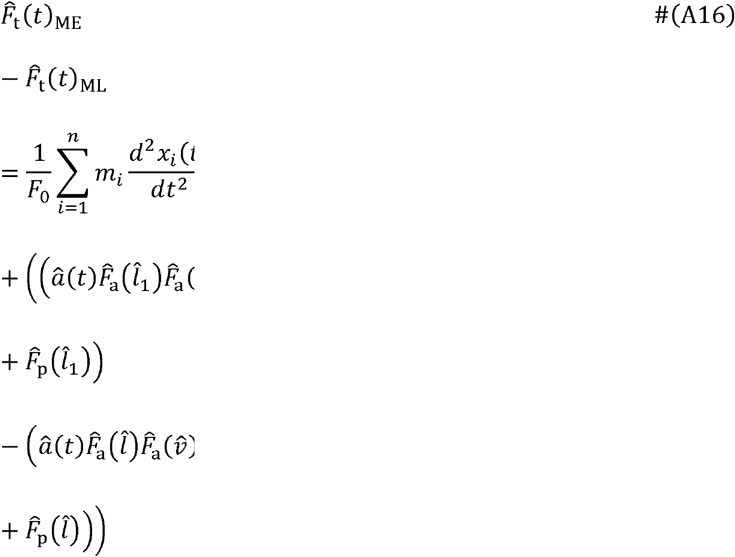

The differences between 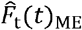 and 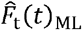 stem from two components. One component is related to 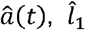 and 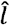. Since velocity is the first derivative of length with respect to time, 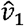 and 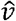 are derived with 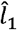, and 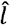, respectively. While 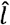 is derived by solving Equations A7–A8 and is unaffected by scale factor, 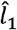 is solved scale factor. Note that â(*t*) contributes to both 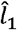, and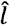 calculations. The other using Equations A9–A10 and is influenced by inertial resistance, thus affected by the component of the difference involves *F*_O_, *m*_*i*_, and *x*_*i*_ (*t*) The point masses *m*_*i*_ scale with scale^3^, *x*_*i*_ (*t*) scale with scale^1^, and *F*_0_ scale with scale^2^, making this component proportional to scale^2^. Consequently, this may explain our findings that RMSD_ME_ is proportional to the range from scale^1.9^ to scale^2^. While the muscle activation level contributes to the RMSD values, its influence might be outweighed by other factors within Equation A16, such as the magnitude of muscle mass and its acceleration.

## Supplementary Information

### Supplementary Figures

**Fig. S1.**
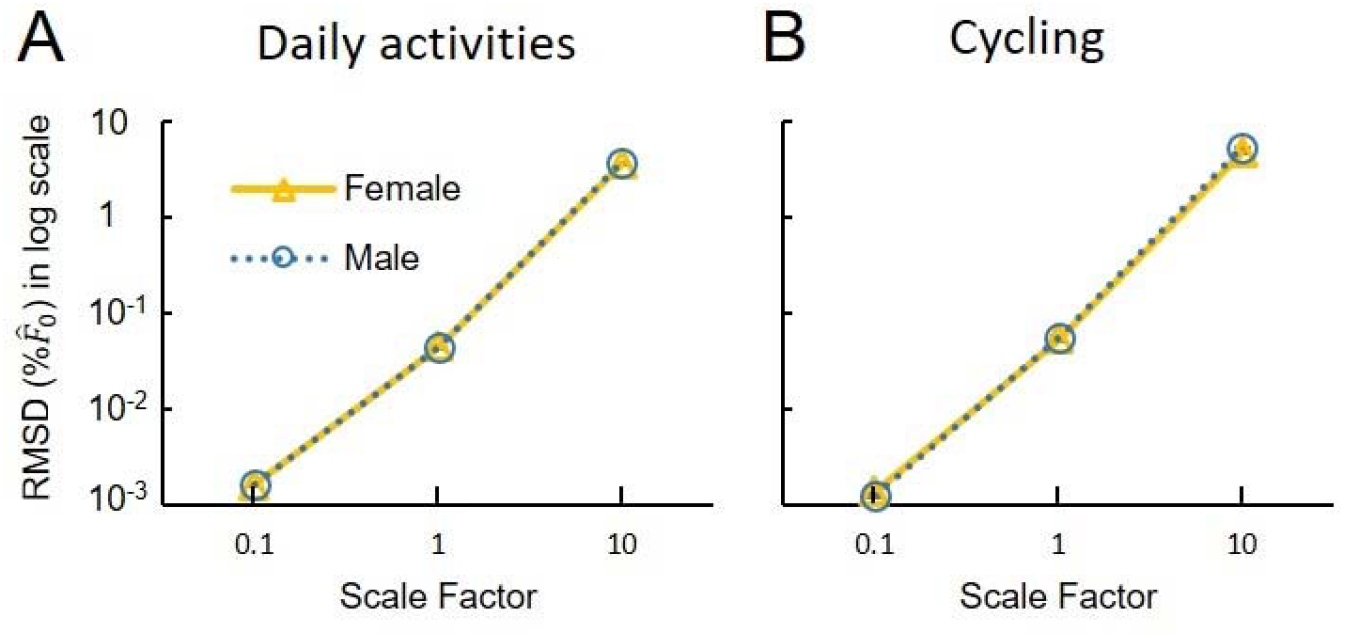
The effect of sex on differences in model-predicted forces across scaled muscle sizes. Each plotted point denotes the average of log-transformed RMSD values across participants, muscles, and tasks for a specific scale and sex. The standard error associated with each mean value is small enough to be encompassed by the mean’s data point. The RMSD values of both sexes overlap nearly completely. ANOVA showed no significant effect of sex on the log-transformed RMSD values across scales.

### Supplementary Tables

**Table S1.** Characteristics of the 20 participants tested. To be included in the study, participants must be adults with normal lower limb function who can independently perform walking, running, hopping, and sit-to-stand movements. People with lower extremity musculoskeletal injuries, neuromuscular diseases, or other systemic diseases affecting their ability to perform the required tasks were excluded.

| Participant | Age (y) | Weight (kg) | Height (m) | Sex | Ethnicity |
| --- | --- | --- | --- | --- | --- |
| 1 | 25 | 54 | 1.75 | F | White |
| 2 | 41 | 66 | 1.67 | M | Asian |
| 3 | 32 | 56 | 1.56 | F | White |
| 4 | 22 | 84 | 1.86 | M | Asian |
| 5 | 26 | 72 | 1.65 | F | Asian |
| 6 | 23 | 77 | 1.78 | M | Middle East |
| 7 | 24 | 47 | 1.55 | F | Asian |
| 8 | 24 | 58 | 1.63 | F | Asian |
| 9 | 27 | 62 | 1.77 | F | Asian |
| 10 | 40 | 72 | 1.73 | M | White |
| 11 | 32 | 72 | 1.71 | M | White |
| 12 | 27 | 65 | 1.63 | F | Asian |
| 13 | 33 | 79 | 1.73 | M | Asian |
| 14 | 41 | 74 | 1.73 | M | Asian |
| 15 | 41 | 75 | 1.93 | M | White |
| 16 | 22 | 77 | 1.83 | M | Asian |
| 17 | 67 | 68 | 1.67 | M | Asian |
| 18 | 40 | 57 | 1.6 | F | Asian |
| 19 | 40 | 57 | 1.63 | F | Asian |
| 20 | 33 | 56 | 1.61 | F | Asian |

